# Parallel genetic screens identify nuclear envelope homeostasis as a key determinant of telomere entanglement resolution in fission yeast

**DOI:** 10.1101/2023.12.26.573359

**Authors:** Rishi Kumar Nageshan, Nevan Krogan, Julia Promisel Cooper

**Author notes:** co-corresponding authors Email: Rishi Kumar Nageshan Nevan Krogan Julia Promisel Cooper.

## Abstract

In fission yeast lacking the telomere binding protein, Taz1, replication stalls at telomeres, triggering deleterious downstream events. Strand invasion from one *taz1Δ* telomeric stalled fork to another on a separate (non-sister) chromosome leads to telomere entanglements, which are resolved in mitosis at 32°C; however, entanglement resolution fails at ≤20°C, leading to cold-specific cell lethality. Previously, we found that loss of the mitotic function of Rif1, a conserved DNA replication and repair factor, suppresses cold sensitivity by promoting resolution of entanglements without affecting entanglement formation. To understand the underlying pathways of mitotic entanglement resolution, we performed a series of genomewide synthetic genetic array screens to generate a comprehensive list of genetic interactors of *taz1*Δ and *rif1*Δ. We modified a previously described screening method to ensure that the queried cells were kept in log phase growth. In addition to recapitulating previously identified genetic interactions, we find that loss of genes encoding components of nuclear pore complexes (NPCs) promotes telomere disentanglement and suppresses *taz1Δ* cold sensitivity; we attribute this to more rapid anaphase midregion nuclear envelope (NE) breakdown in the absence of these NPC components. Moreover, loss of genes involved in lipid metabolism reverses the ability of *rif1*+ deletion to suppress *taz1Δ* cold sensitivity, again pinpointing NE modulation. A *rif1*+ separation-of-function mutant that specifically loses Rif1’s mitotic functions yields similar genetic interactions. Genes promoting membrane fluidity were enriched in a parallel *taz1+* synthetic lethal screen at permissive temperature, cementing the idea that the cold specificity of *taz1Δ* lethality stems from altered NE homeostasis.

## Introduction

Telomeric sequences constrain replisome passage (Bochman et al. 2012; Miller et al. 2006). In fission yeast, binding of the conserved double strand telomere binding protein Taz1 promotes replisome passage; an analogous function in promoting telomeric semi-conservative replication is performed by mammalian TRF1, a Taz1 ortholog (Sfeir et al. 2009). In the absence of Taz1, stalled telomeric replication forks accumulate and are processed by the RecQ type DNA helicase, Rqh1, rendering them incompetent to resume replication (Rog et al. 2009). These irreversibly stalled forks occur at multiple telomeres, resulting in strand invasions between them that result in non-sister chromosome entanglements (Figure 1A) (Miller and Cooper 2003; Miller et al. 2006; Zaaijer et al. 2016). Cycling *taz1*Δ cells maintained at temperatures at or above 25°C (≥25°C is used as permissive temperature herein) successfully resolve these entanglements (Miller and Cooper 2003). However, when grown at ≤20°C (hereafter referred to as ‘cold’), entanglement resolution fails, causing cold sensitivity (hereafter referred to as ‘c/s’) (Figure 1B).

**Figure 1:**
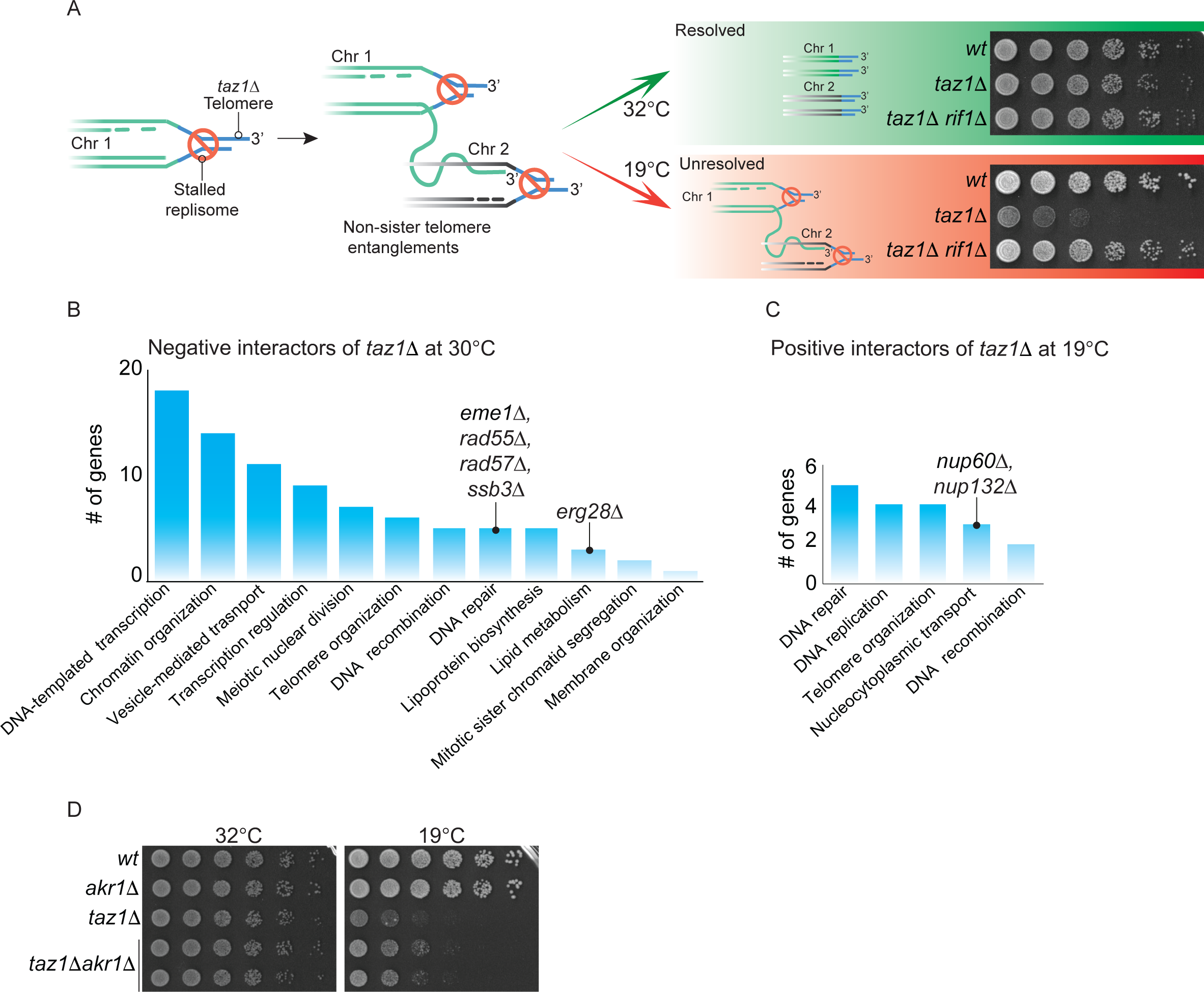
Genes involved in telomere entanglement formation, resolution and cold sensitivity. ***A***, Representation of telomere entanglement formation in absence of Taz1. The green and red arrows show the fates of these entanglements: they are resolved when the cells are grown at 32°C, but resolution fails when cells are grown at 19 °C, causing c/s. On the right, 5-fold serial dilution of logarithamically grown cells incubated at 32°C (2 days) or 19 °C (7 days) are shown. Rif1 inhibits the resolution of *taz1Δ* telomere entanglements in the cold. ***B***, Graph of number of genes from the respective GO groups that show negative genetic interactions with *taz1*Δ at 30°C. ***C***, Graph of number of genes from the respective GO groups that show synthetic lethality with *taz1*Δ cells at 19°C. Note: only selected groups are represented here; an unedited GO tag enrichment list is provided in Table S3 and S4 for ***B*** and ***C***, respectively. ***D***, 5-fold serial dilution of logarithmically growing cells incubated at 32°C (2 days) or 19 °C (7 days).

To understand the mechanism of entanglement formation, we previously performed an overexpression screen to identify suppressors of *taz1*Δ c/s (Rog et al. 2009). Overexpression of the SUMO deconjugation enzyme Ulp1 or the G1/S transcription factor Cdc10, which controls Ulp1 expression, suppresses *taz1*Δ c/s. These suppressors pointed us towards the action of sumoylated Rqh1, which processes stalled telomeric replication forks to prevent their restart, instead channeling them into entanglement formation in S-phase (Rog et al. 2009). The above-described screen left open the question of what factors control entanglement resolution.

The conserved DNA replication and repair protein Rif1 promotes *taz1*Δ c/s (Figure 1B), as *rif1Δ* confers strong suppression of *taz1Δ* c/s (Miller and Cooper 2003; Zaaijer et al. 2016). The loss of Rif1 does not impact the stalling of telomeric replisomes or formation of entanglements, but rather promotes disentanglement. The development of a *rif1*+ separation-of-function (*S-rif1+*) allele allowed us to delineate Rif1’s anaphase function specifically as relevant to the resolution of entanglements and suppression of c/s (Zaaijer et al. 2016). To address how Rif1 controls the resolution of entanglements as well as to compile a full array of genes that determine *taz1Δ* viability, we performed a series of genomewide synthetic genetic array screens to map genetic interactions with *taz1*+ and *rif1*+. To avoid unrelated issues regarding return from stationary phase growth to mitotic growth in the absence of Taz1, we developed a protocol to keep the cells in log phase throughout all robotic pinning steps. The resulting hits were sorted to select gene deletions that 1) are synthetic sick/lethal with *taz1Δ* alone at the permissive temperature, 2) suppress *taz1Δ* c/s and 3) reverse *taz1Δ* c/s suppression by *rif1Δ* or *S-rif1+*. The set of genes thus obtained will inform, respectively, on 1) processes that restrain entanglement formation and/or recapitulate properties conferred by cold temperature, 2) pathways that impact either formation or resolution of entanglements, and 3) pathways that specifically control *taz1Δ* telomere disentanglement in mitosis. Our screening results revealed not only interacting genes from the DNA replication-repair and chromatin organization ontology groups, but also genes involved in nuclear pore complex (NPC) and lipid homeostasis. Collectively, we provide a comprehensive genetic interaction map of the regulators of telomere entanglement resolution and the final steps of mitosis, in which timely nuclear envelope (NE) remodeling plays a surprisingly crucial role.

## Materials and Methods

### Media

Media comprised YES broth (Sunrise #2011-500), YES agar (Sunrise #2012-500), PMG (Sunrise #2060-500). Media and growth conditions were as previously described (Moreno et al. 1991). Strains used are listed in Table S1.

### Strain construction

Gene deletions were generated as described previously and used to create further strains through crossing, sporulation and selection (Bahler et al. 1998).

## Synthetic genetic array screening and analysis

We modified the previously described protocol (Roguev et al. 2007; Roguev et al. 2018) to generate fission yeast genetic interactions maps. Most of the previous genetic screens were performed on stationary phase cells with an end point assay, querying viability or sensitivity to environmental challenges. To avoid enrichment for genes that impact cell fitness in or following stationary phase, cells were subjected to a modified protocol. Logarithmically growing *wt* liquid cultures show ∼15-20% septating cells (septation index). We recreated a similar septation index profile by repeated robotic pinning at time intervals short enough to avoid stationary phase, and subjected log phase cells to growth at 19°C (Figure S1A).

Single or double mutants of *taz1Δ, rif1Δ,* and/or *S-rif1+* were generated in the PEM-2 strain background as described previously (Roguev et al. 2018). We confirmed that the *taz1Δ* c/s and suppression of c/s by *rif1Δ* are unchanged in the PEM-2 background (Figure S1B). Our screening protocol is outlined in Figure S1C. Briefly, the query strains were crossed with the Bioneer *S. pombe* gene deletion library. Double or triple mutant haploids derived from *taz1Δ, rif1Δ, S-rif+*, *taz1Δrif1Δ, or taz1ΔS-rif1+* were selected. The double mutants were selected on cycloheximide, hygromycin and G418 and the triple mutants on cycloheximide, hygromycin, G418 and nourseothricin containing media. These selected clones were plated robotically as described earlier with two additional rounds of repetitive stamping and a reduced incubation time (as outlined in Figure S1C) at 30°C on YES plates. The final stamping was on two sets of plates. One set was incubated at 30°C and the other at 19°C; both lacked antibiotic selection, to minimize any impact on cell fitness. Note that dilution assays to validate the screening results are performed at 32°C, as majority of our previous results have been documented at this temperature; moreover, as mentioned above, at temperatures ≥25°C, *taz1Δ* cells do not show any growth defects. The screen was performed in technical triplicates. After 3 days of growth the plates were imaged and growth (colony size) was measured as described earlier (Roguev et al. 2018). Growth of each genotype was normalized to the median growth on a given plate referred as Normalized Growth (NG). The NG at 30 °C was directly used as a measure of genetic interactions between single mutant and genes from the library (as with *taz1Δ* in Figure 1B). The NG ratio (NGR) at 19°C and 30°C was calculated for each genotype as a measure of genetic interactions at 19°C; all unedited normalized growth ratios are listed in (Table S2). Outliers were determined by the 2-standard deviation method for high stringency hits (followed for *taz1Δ* interactions, table S3 and S4). In addition, the top 25% percentile of genetic interactors (with *rif1Δ,S-rif1+, taz1Δrif1Δ, and taz1ΔS-rif1+*; table S2 and S5) were further analyzed to select specific hits that restored the anaphase role of *rif1*+ in promoting *taz1Δ* c/s. The top 25^th^ percentile was chosen to avoid losing any relevant hits, as *rif1Δ* emerged as a *taz1Δ* suppressor in this 25^th^ percentile. Gene ontology (GO) enrichments were determined using www.pombase.org (biological process slim) inbuilt GO ontology tags.

## Results and Discussion

To investigate the mechanisms that control *taz1Δ* entanglement resolution, we performed a series of parallel screens in which the *S. pombe* gene deletion library was crossed with *taz1Δ*, *rif1Δ* and *taz1Δrif1Δ* cells to identify those gene deletions that rescue or exacerbate *taz1Δ* c/s or reverse the suppression of c/s conferred by *rif1+* deletion. We also employed the *S-rif1+* allele, which acts as a null specifically for Rif1’s mitotic functions, in parallel screens to distinguish those factors that act at S- versus M-phase (Zaaijer et al. 2016). Based on our previous work, we expected a number of genetic interactions with *taz1Δ*. For instance, our previous studies showed that *rap1+* deletion confers synthetic sickness with *taz1Δ* at 19°C, while *ulp2Δ* and *rif1Δ* are suppressors of *taz1Δ* c/s; these interactions appeared in our screens (Table S2), validating their efficacy (Miller and Cooper 2003).

## Replication fork processing factors are required for *taz1*Δ cell viability

To identify the players that control *taz1*Δ cell viability at permissive temperature, we ascertained specific genetic interactions with *taz1*Δ by filtering out any hits that show similar interaction with *rif1*Δ (Table S3). GO analysis of the filtered hits revealed genes involved in transcription as the top hits, followed by factors controlling chromatin organization. These factors could change the chromatin landscape of *taz1*Δ telomeres or may act indirectly via controlling expression of other genes (Figure 1B).

The stalled replication forks at *taz1Δ* telomeres are processed by DNA replication and repair factors such as Rqh1(Rog et al. 2009), which promotes the aberrant processing of these forks to form entanglements. The gene encoding Eme1, a component of an endonuclease complex that promotes homologous crossovers, is required for full *taz1*Δ cell viability, as are the DNA replication/repair factors Rad55, Rad57 and the Replication Protein complex A subunit Ssb3. It will be important to dissect whether these factors act at the level of entanglement generation, mitotic entanglement resolution, or suppression of nonhomologous end-joining in G2 cells (Ferreira and Cooper 2001; 2004).

## Loss of nuclear pore complex components suppresses *taz1***Δ** c/s

Several of the strongest suppressors of *taz1*Δ c/s encode components of NPCs (Figure 1C and Table S4). We have validated the positive genetic interactions with *nup60*Δ and *nup132*Δ and performed an extensive functional analysis (Nageshan et al. 2023). This analysis pinpointed the functions of Nup60 and Nup132 in a local NE breakdown event as crucial to the resolution of *taz1Δ* telomeric entanglements. Briefly, fission yeast undergo NE breakdown in a temporally and spatially restricted manner, at the anaphase midregion (Dey et al. 2020; Exposito-Serrano et al. 2020), across which any persisting chromosome entanglements are stretched. In a *wt* cell this NE breakdown exposes the mitotic spindle to cytosol, which is essential for spindle disassembly and a successful nuclear divison. The midregion harbors NPCs that are sequentially dismantled, leading to the local NE breakdown. Loss of NPC components hastens this process, ensuring a fragile midregion (Nageshan et al. 2023). Along with our analysis of key hits in the parallel screen for reversal of suppression by *rif1Δ* (see below), our data show that advancing NE breakdown promotes telomere entanglement resolution.

## Lipid membrane homeostasis regulates entanglement resolution

Genes that control lipid biosynthesis and membrane homeostasis (Figure 2B) emerged as prominent genetic interactors with *taz1Δ*. In mammalian cells, cholesterol controls membrane fluidity, with higher levels of cholesterol generally conferring greater fluidity; ergosterol carries out these functions in the membranes of fungi. The gene encoding Erg28, which is involved in ergosterol biosynthesis, is required for *taz1*Δ cell survival even at permissive temperature (30°C; Figure 1B and Table S3). Budding yeast cells lacking Erg28 have been shown to harbor a three-fold reduction in ergosterol along with accumulation of 3-keto sterols and carboxylic acid sterols and therefore, display reduced membrane fluidity (Gachotte et al. 2001; Mo and Bard 2005). This genetic interaction suggests that membrane fluidity promotes resolution of telomere entanglements and is consistent with our observation that treatment with membrane fluidizing agents suppresses *taz1Δ* c/s (Nageshan et al. 2023). We hypothesize that enhanced NE rigidity mimics cold temperatures, which increase phospholipid membrane rigidity. In either cold conditions or at 30°C in the absence of Erg28, NE fluidity is reduced, confounding entanglement resolution by inhibiting anaphase midregion NE breakdown.

**Figure 2:**
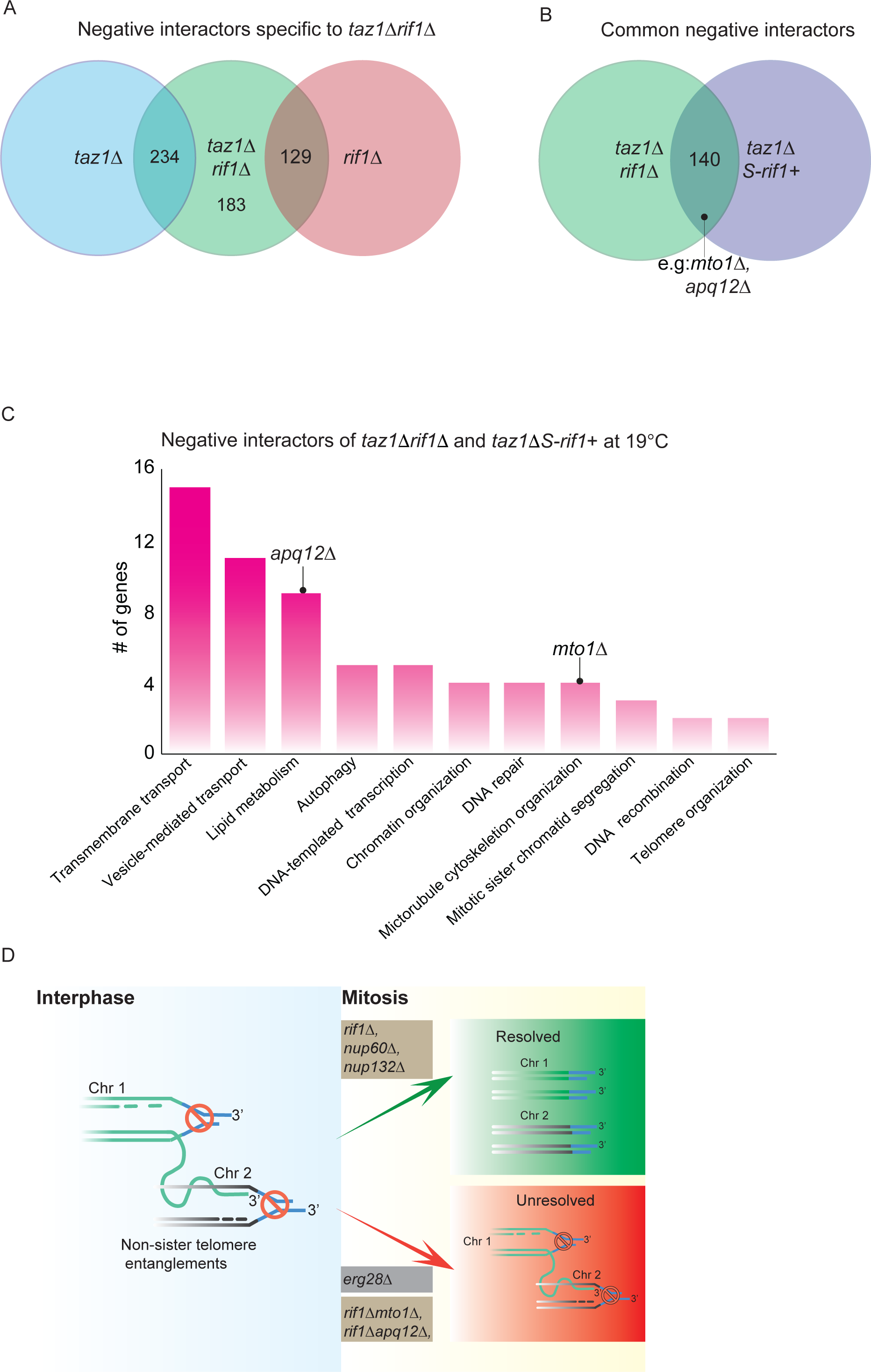
Genes that revert the ability of *rif1*Δ to suppress *taz1Δ* anaphase phenotype. ***A***, Venn diagram of the genes that show negative interactions with *taz1*Δ*rif1*Δ, specifically in the top 25th percentile with respect to the strength of the reversion. ***B***, Venn diagram of interactors common to *taz1Δrif1Δ* and *taz1ΔS-rif1*+. The common genes specifically restore the anaphase Rif1 function that is relevant to *taz1Δ* telomere entanglement resolution. ***C***, Graph of number of genes from selected GO groups that show negative interactions with *taz1*Δ*rif1*Δ and *taz1*ΔS-*rif1*+ cells at 19°C. An unedited summary of the GO term analysis is provided in Table S5. ***D***, Representation of telomere entanglement as in Figure 1A, with the addition of genetic interactions observed herein. Gene deletions that enable resolution are above the green arrow, while those that hinder resolution are at the red arrow. Genotypes that are highlighted in beige represent interactions at 19°C and those highlighted in grey represent interactions at 32 °C.

In contrast to the synthetic sickness between *erg28Δ* and *taz1Δ*, deletion of the gene encoding the Akr1 palmitoylacyltransferase confers a mild suppression of *taz1Δ* c/s. A recent report showed that Akr1 localizes to the nuclear periphery as well as the cell tips and is required for fusion of the two NEs during meiosis (Pham et al. 2023); in the absence of Akr1, karyogamy fails. We generated a new double mutant by deleting *akr1*+ in the *taz1Δ* background to avoid the need to undergo meiosis, and confirmed c/s suppression of *taz1Δ* by *ark1Δ* (Figure 1D and Table S2). Though we do not yet know the relevant palmitoylation targets, it is tempting to speculate that Rif1 might be a target. Previous reports have demonstrated that budding yeast Rif1 is targeted to the inner NE via palmitoylation (Fontana et al. 2019; Fox and Gartenberg 2012; Park et al. 2011); such targeting facilitates the ability of Rif1 to direct DNA double strand break repair away from resection-dependent homologous recombination and towards nonhomologous end-joining (Fontana et al. 2019), neither of which control *taz1Δ* telomere entanglements (Miller and Cooper 2003; Rog et al. 2009). Hence, while these previous studies focused on Rif1’s role in S-phase or interphase, palmitoylation may also play a role in Rif1’s mitotic localization and actions.

## Loss of genes involved in lipid metabolism restores Rif1’s anaphase function in regulating telomere entanglement resolution

To delineate the molecular mechanisms of Rif1’s anaphase function, we screened for gene deletions that specifically revert the suppression of *taz1Δ* c/s by *rif1*Δ (Figure 3A). We also carried out the analogous screen for reversion of *taz1Δ* c/s in a *S-rif1*+ background in which Rif1’s S-phase functions are retained while its mitotic function is lost (Zaaijer et al. 2016). GO analysis of the gene deletions that reverted both *rif1*Δ and *S-rif1+* phenotypes show lipid metabolism as one of the most prominent ontology groups (Figure 2C and Table S5). Among these, *apq12+* loss reverted Rif1’s mitotic function, conferring c/s to *taz1Δrif1Δ* cells without affecting the c/s of single *taz1Δ* mutants. Budding yeast Apq12 is a NE and endoplasmic reticulum protein whose loss leads to defects in NPC assembly stemming from deregulation of NE lipid homeostasis (Scarcelli et al. 2007; Schneiter and Cole 2010). The defects in NPC assembly in *apq12Δ* cells are averted by treatment with low concentrations of the membrane fluidizing agent benzyl alcohol (BA). In fission yeast, *apq12*Δ cells have likewise been shown to suffer defects in NE homeostasis (Tamm et al. 2011). Altered NE dynamics in the absence of Apq12 would be compatible with a model in which more rapid NPC disassembly, afforded by Rif1 loss, is counteracted by severe levels of NE rigidity in *apq12Δ* cells at 19°C, delaying anaphase midregion NE breakdown and reverting *taz1Δ* viability.

Our analyses also pinpointed a gene not previously implicated in NE homeostasis, *mto1+*, as a key player in the dynamics of anaphase midregion NE breakdown. Mto1 emerged as a gene product whose loss strongly reverts the suppression of *taz1Δ* c/s by *rif1+* deletion. We confirmed and analyzed this genetic interaction (Nageshan et al. 2023). Mto1 has been known as a cytoplasmic microtubule organizing protein whose loss leads to the disappearance of several classes of cytoplasmic microtubules, generating high concentrations of free tubulin and in turn, spindle persistence (within the nucleus) (Sawin et al. 2004). However, we found that *mto1Δ* cells also suffer delayed anaphase midregion NE breakdown, possibly stemming from the loss of cytoplasmic MTs that physically prod the anaphase midregion NE and stimulate its disassembly (Nageshan et al. 2023). Our analysis included separation of Mto1’s spindle persistence and NE breakdown-promoting functions, and demonstrated that Mto1 promotes entanglement resolution via advancement of anaphase midregion NE breakdown.

## Vesicle-mediated transport process regulates telomere disentanglement

Gene deletions that have been classified to function in vesicle-mediated transport show negative genetic interactions with *tazΔ* at 30°C (Figure 1C) and with *taz1Δrif1Δ* at 19°C (Figure 2C). Genes (*vsp26+*, *sft1*+, *tlg2*+) encoding proteins that are involved in transport of vesicles between the ER and Golgi apparatus, along with a gene (*dot2*+) encoding an ESCRTII complex subunit, were required for *taz1Δ* viability at 30°C. Similarly, loss of *sft2+*, *psg1+* or SPAC9ES.04 reversed suppression of *taz1Δ* c/s by *rif1Δ*. A recent report presents a comprehensive physical interaction screen between fission yeast NPCs and the inner NE proteins Ima1 and Lem2, using the yeast two-hybrid approach (Varberg et al. 2020). This study found that many of the proteins encoded by the above-mentioned genes interact with inner NE proteins; for instance, Ima1 interacts with Sft1, Sft2, Psg1, and SPAC9ES.04p. Additionally, Cut11, a component of both the NPC and the spindle pole body, shows interactions with this set of gene products. These interactions highlight a functional interplay between NE dynamics, genes classified as involved in vesicle transport, and *taz1Δ* telomere disentanglement. These observations herald an exploration of how chromosome entanglements in the anaphase midregion communicate with the NE breakdown machinery, and how membrane sorting/repair proteins participate in this process.

Collectively, the synthetic genetic array screens hold a wealth of information, prominently suggesting unforeseen links between DNA processing events and regulators of NE fluidity and NPC disassembly (Figure 2D). The synthetic sickness between *erg28Δ* and *taz1Δ* suggests that membrane rigidity is a key factor in mediating the cold-specificity of *taz1Δ* lethality. Additional genes whose functions have been implicated in NE fluidity or NPC dismantlement-induced anaphase NE breakdown play crucial roles in *taz1Δ* telomere entanglement. Moreover, in-depth analyses of genes classified in GO groups seemingly unrelated to NE breakdown may uncover roles in this process, as has been the case for Mto1 (Nageshan et al. 2023). A comprehensive analysis of the DNA repair/replication/recombination genes identified will also be crucial for dissecting the mechanisms that make NE breakdown, and therefore cytoplasmic exposure of telomere entanglements, so crucial for their resolution.

## Acknowledgements

We thank our lab members for discussions and valuable input, particularly Haitong Hou (now at Jiangnan University) who provided critical comments that helped in designing the analysis pipeline. This work was supported by the NCI and the University of Colorado School of Medicine.

## Author contributions

RKN performed all the experiments. NK hosted RKN in his lab to perform the synthetic genetic array screens. RKN and JPC conceived the study, designed and interpreted the experiments, and wrote the manuscript. The authors declare no conflict of interest.

## Supplementary information

**Figure S1:**
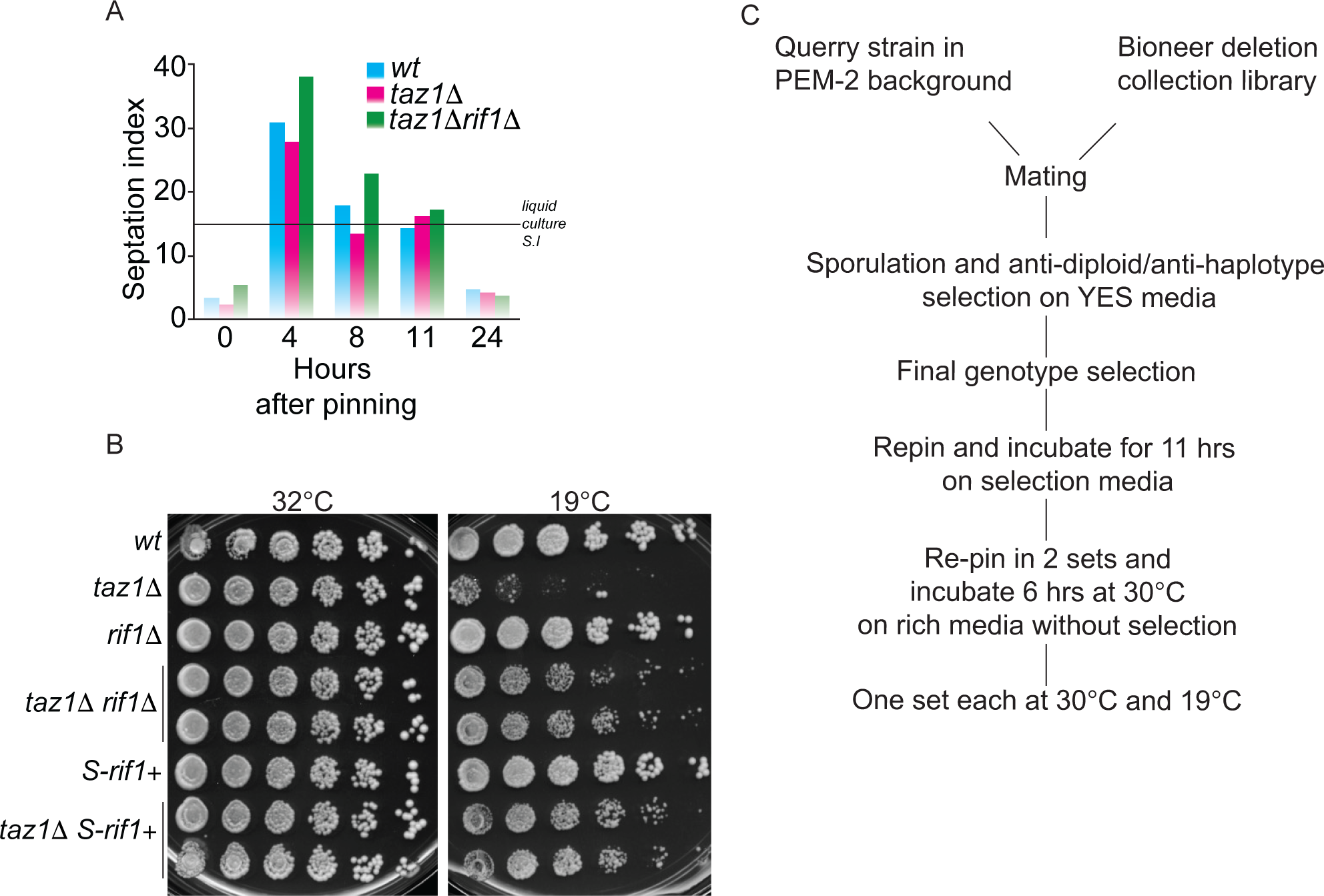
***A***, Standardization of pinning timing to ensure that queried cells are in log phase. *wt* cells from a lawn grown for 16 hours were manually pinned onto a fresh YES plate and incubated at 32°C. Septation index was determined by calcofluor staining at the represented time points after pinning. In parallel, a *wt* culture was maintained in log phase (0.5 OD) and the septation index was calculated. The observed septation index (∼15%) is indicated as a dotted line on the graph. ***B***, 5-fold serial dilution of logarithamically grown cells incubated at 32°C (2 days) or 19 °C (7 days). All the strains were created in PEM-2 strain background. ***C***, Summary of the protocol used for the genetic screen. The septation index of a *wt* colony from each plate was manually checked (not shown here) before the plates were moved to their respective temperatures.

**Tables: All tables can be found at the following link:**

(https://figshare.com/s/7a769b7f1f6248b27058)

**Table S1:** List of strains used in the current study.

**Table S2**: Unedited list of the top 25th percentile interaction strength genes that show interactions with the indicated query strains and corresponding NGR values at 19°C and 30°C.

**Table S3:** List of genes that show negative genetic interaction with *taz1Δ*. The first sheet shows systematic names along with respective NG values (see Methods). The second sheet shows the gene information for the systematic names in sheet 1. Sheet 3 shows the GO tags for sheet 1. Sheet 4 is a summary of sheet 3, indicating the number of genes corresponding to each GO tag. The same format is followed for Tables 4 and 5 as well.

**Table S4:** Unedited list of genes that show positive genetic interaction with *taz1Δ.* The sheet format follows that of Table 3.

**Table S5:** List of genes whose deletion reverses the anaphase function of *rif1+* in suppresses *taz1Δ* c/s. The sheet format follows that of Table 3.

